# Enhancing the cryopreservation of *ex vivo* 3D tumor models using vitrification strategies

**DOI:** 10.1101/2025.10.01.679784

**Authors:** Tommy Brasseur, Gabriel Pagé, Amélie St-Georges-Robillard, Anne-Marie Mes-Masson, Thomas Gervais

## Abstract

Microdissected tumor tissue explants (MDTs) are promising ex vivo models for oncology research but remain limited by poor preservation and the rapid loss of viability following resection. Here, we systematically optimized MDT cryopreservation using prostate-derived 49F and ovarian TOV112D tumor models. Before cryopreservation, we evaluated the effects of antioxidant supplementation, pre-cooling, and a short recovery period. MDTs were then preserved by either conventional slow-freezing or vitrification using ultra-rapid cooling. Following thawing, tissue morphology, apoptosis, and proliferative capacity were assessed relative to fresh controls. Vitrification improved morphological preservation and reduced apoptosis compared with slow-freezing, although recovery of proliferation differed between tumor models. Antioxidant supplementation enhanced post-thaw proliferation at optimal concentrations but induced toxicity at higher doses. Pre-cooling and a short pre-cryopreservation recovery period further improved post-thaw outcomes. Combining these parameters produced an optimized vitrification protocol that preserved up to 98% of the proliferative capacity of 49F MDTs and 69% of that of TOV112D MDTs relative to fresh controls. These findings establish optimized vitrification as a reproducible, high-yield strategy for preserving MDTs for downstream ex vivo oncology applications.

## I. Introduction

Submillimeter-scale tumor explant models are increasingly regarded as powerful tools in biomedical research due to their ability to preserve the native three-dimensional architecture, cellular heterogeneity, and microenvironmental context of *in vivo* tumors[1] [2], [3]. These models have been widely used for applications ranging from drug screening to the study of tumor biology and personalized medicine[4], [5], [6], [7]. However, despite their advantages, maintaining the viability and functional integrity of tumor explants ex vivo remains a significant challenge. Their limited size, lack of vascularization, and sensitivity to environmental conditions contribute to rapid degradation and cell death if not handled under tightly controlled conditions.

To address these limitations, reducing tissue dimensions to the submillimeter scale has emerged as an effective strategy to improve viability. When tumor tissue is microdissected into sufficiently small fragments, passive oxygen diffusion can replace vascular transport, enabling sustained survival ex vivo for several days and, in some cases, up to two weeks [8]. This approach also allows for the generation of multiple microdissected tissue explants (MDTs) from a single tumor sample, which is particularly advantageous given the limited availability of patient-derived material [9]. Such scalability is critical for enabling parallelized testing of therapeutic conditions in the context of personalized medicine.

Microfluidic chips further enhance the utility of these explants by enabling precise control over their microenvironment while facilitating their manipulation at small scales. These systems allow for stable positioning of explants, controlled delivery of nutrients and drugs, and efficient oxygenation, while minimizing reagent consumption. Over the past decade, microfluidic approaches have been increasingly applied to tissue explant culture, including organ-on-chip systems and platforms for functional tissue analysis[10], [11]. In this context, the combination of MDTs and microfluidic technologies provides a powerful framework for high-throughput and physiologically relevant ex vivo experimentation.

Despite these advances, the broader clinical and research adoption of explant-based platforms remains limited by the inherent instability of fresh tissue. Tumor explants are highly sensitive to handling conditions, including temperature fluctuations and delays between resection and processing, which can significantly impact viability. Furthermore, the lack of reliable preservation methods prevents long-term storage and centralized processing, thereby limiting opportunities for retrospective studies and large-scale biobanking. In practice, variability in surgical schedules further complicates tissue collection, as samples obtained outside standard working hours are often discarded due to the inability to process them in time [12], [13], [14]. Developing robust preservation strategies is therefore essential to expand the accessibility and impact of explant-based models.

Cryopreservation represents a promising solution to these challenges, and recent advances in microfluidic technologies have further expanded opportunities to improve cryoprotective agent delivery, vitrification, and post-thaw tissue handling[15]. Broadly, cryopreservation techniques can be divided into two main categories: slow freezing and vitrification. Slow freezing involves a gradual decrease in temperature (typically 1 °C per minute) in the presence of low concentrations of cryoprotective agents (CPAs), such as dimethyl sulfoxide (DMSO). While this approach is well-established for cell suspensions, it is poorly suited for three-dimensional tissue explants [12]. The limited diffusion of CPAs into the tissue core can result in intracellular ice formation and structural damage, even under controlled cooling conditions. Increasing CPA concentration may improve penetration, but this approach is constrained by cytotoxicity, particularly for metabolically active tissues [16].

On the other hand, vitrification avoids ice formation by transitioning the solution into an amorphous glass-like state through the use of high CPA concentrations and ultrarapid cooling, typically achieved via liquid nitrogen immersion [17], [18], [19]. This method has the potential to better preserve tissue architecture while minimizing ice-induced damage. However, vitrification remains underexplored for complex tissue explants, in part due to challenges in achieving consistent post-thaw viability [20], [21].

The success of vitrification in other biomedical fields further supports its potential for preserving complex biological systems. In assisted reproductive technologies, vitrification has largely replaced conventional slow freezing as the preferred method for the cryopreservation of oocytes and embryos due to its superior post-thaw survival, reduced ice crystal formation, and improved clinical outcomes[22], [23], [24]. Today, vitrification is considered the clinical standard for embryo and oocyte preservation in *in vitro* fertilization (IVF) programs worldwide. These achievements demonstrate the ability of vitrification to preserve multicellular structures with high functional integrity and suggest that similar benefits could potentially be extended to other complex tissues. However, whether these advantages translate to heterogeneous tumor explants remains largely unexplored.

Recent studies suggest that supplementing cryopreservation protocols with antioxidants may improve outcomes by mitigating oxidative stress associated with freezing and thawing. In reproductive medicine, antioxidant combinations such as acetyl-L-carnitine, N-acetyl-L-cysteine, and α-lipoic acid (A3) have been shown to enhance embryo viability and developmental potential under stress conditions. Similarly, individual antioxidants such as ascorbate have demonstrated protective effects during cryopreservation[25], [26].

Another important but often overlooked factor influencing cryopreservation success is the mechanical stress induced during tissue preparation. Unlike cell suspensions, which are processed under relatively controlled conditions, tissue explants are obtained through surgical resection and subsequent microdissection, processes that introduce mechanical disruption and potential damage. Previous work has shown that short-term culture on microfluidic chips can partially restore viability and promote proliferation of MDTs following dissection. This suggests that allowing a recovery period prior to cryopreservation may improve outcomes by reducing stress-related damage, although such approaches currently require specialized infrastructure.

In this study, we systematically compared slow freezing and vitrification for the cryopreservation of microdissected tumor explants derived from two cancer models: TOV112D ovarian tumors and modified 49F prostate tumors. The vitrification protocol investigated in this study was adapted from clinically established protocols developed for assisted reproductive technologies, with modifications to accommodate the unique characteristics of tumor tissue explants. We further evaluated whether antioxidant supplementation, temperature modulation, and a pre-cryopreservation recovery period could improve post-thaw tissue preservation. Morphology, apoptosis, and proliferative capacity were assessed to identify the combination of conditions providing the most reproducible and effective preservation. Our objective was to establish a simple, high-yield cryopreservation strategy that could improve the accessibility and translational utility of explant-based platforms for cancer research and personalized medicine.

## II. Material and methods

### 2.1 : Microfluidic chip

The microfluidic chip used in this study was originally described by Simeone et al. [27] and purchased from MISO Chip (Montreal, Canada). The device is fabricated in polydimethylsiloxane (PDMS) and consists of four parallel channels, each containing eight cubic wells of 700 µm width. MDTs were introduced into the chip by pipetting and allowed to settle into the wells by sedimentation, where they remained throughout the experiment [10]. Nylon cylinders were placed at the inlets to generate hydrostatic pressure for medium exchange and treatment administration.

Prior to use, microfluidic devices were passivated and sterilized following previously published protocols [23], [24]. Briefly, channels and wells were first treated with a triblock copolymer (Pluronic® F-108, 542342, Millipore- Sigma, St. Louis, USA) to reduce cell adhesion to PDMS surfaces. Devices were then sterilized using 70% ethanol (P016EAAN, Commercial Alcohols) and rinsed with sterile Hank’s Balanced Salt Solution (HBSS, 311-425-CL, Wisent, Canada). Prepared chips were filled with HBSS and stored at 4 °C until use.

### 2.2 : Cell line and xenograft generation

Human ovarian cancer cells TOV112D (RRID: CVCL_3612) were cultured in OSE medium (316-030-CL, Wisent, Canada) supplemented with 10% fetal bovine serum (FBS, 098-150, Wisent), 0.6 mg L^-1^ amphotericin B (450-105- QL, Wisent), and 55 mg L^-1^ gentamicin (450-135-XL, Wisent). Human prostate cancer cells 49F (RRID: CVCL_RW53), obtained from A. Zoubeidi (University of British Columbia, Canada) were cultured in RPMI 1640 medium (350-000-CL, Wisent) supplemented with the same additives. All cells were maintained at 37 °C in a humidified incubator with 5% CO_2_.

For xenograft generation, cells were detached using trypsin-EDTA (0.25%, 25200056, Thermo Fisher Scientific, Waltham, USA) and resuspended in a 1:1 mixture of Matrigel (356237, Corning) and phosphate-buffered saline (PBS, 311-425-CL, Wisent) to obtain a final concentration of 2 10 cells in 400 µL. Cell suspensions were injected subcutaneously into the flank of NRG mice (007799, The Jackson Laboratory, Bar Harbor, ME, USA). Male mice were used for 49F xenografts and female mice for TOV112D xenografts.

All animal procedures were approved by the Institutional Animal Care Committee (IACC) of the Centre de recherche du Centre hospitalier de l’Université de Montréal (CRCHUM) under protocol CIPA C22014AMMs.

### 2.3 : MDT generation, loading, and culture conditions

Microdissected tumor explants (MDTs) were generated from freshly excised xenograft tumors using an adaptation of previously described protocols [8]. Briefly, tumors were first sectioned into 350 µm-slices using a McIlwain tissue chopper (10180, Ted Pella, Redding, CA, USA) and maintained in HBSS supplemented with 10% fetal bovine serum (FBS, 098–150, Wisent), 55 mg L^-1^ gentamicin, and 0.6 mg L^-1^ amphotericin B to preserve tissue viability during processing. Tissue fragments were then further processed into approximately spherical MDTs using a 500 µm biopsy punch (PUN0500, Zivic Instruments, Pittsburgh, USA), resulting in explants of approx. 350-400 µm in diameter. Following microdissection, MDTs were maintained in HBSS supplemented with 55 mg L^-1^ gentamicin, and 0.6 mg L^-1^ only until loading.

MDTs were loaded into microfluidic chips following previously established procedures [8]. For each channel, eight MDTs were collected using a micropipette and introduced into the inlet reservoirs filled with HBSS. Controlled flow was generated by gentle aspiration at the outlet, allowing MDTs to travel through the channel. When an MDT reached the vicinity of a trapping well, flow was temporarily halted (approx. 1-2 s) to allow sedimentation into the well. This process was repeated until all four channels were loaded, resulting in a total of 32 MDTs per chip. After loading, HBSS was replaced with the appropriate culture medium supplemented with 10% FBS, 55 mg L^-1^ gentamicin, and 0.6 mg L^-1^ amphotericin B. Chips were incubated at 37 °C with 5% CO_2_ in a humidified environment to minimize evaporation from the microfluidic channels. Culture medium was renewed every 48 h.

At experimental endpoints, MDTs were fixed in formalin and processed into formalin-fixed paraffin-embedded (FFPE) blocks using histopathological methods adapted for microfluidic devices, as previously described [27].

### 2.4 : recovery culture pre-freezing

To evaluate whether a recovery period following microdissection improved cryopreservation outcomes, MDTs were cultured for one, two, or four days prior to freezing. Recovery was performed either in the microfluidic chip or in conventional Petri dishes. This comparison was designed to assess whether recovery benefits depended on the microfluidic chip or could also be achieved using standard culture conditions. Fresh, non-cryopreserved MDTs were cultured in parallel for the same duration and served as controls.

### 2.5 : Cryopreservation and thawing solutions

Cryopreservation experiments were performed on groups of approximately 80 MDTs per condition. For vitrification protocols, MDTs were transferred into 2 mL cryotubes and exposed to cryoprotectant (CPA) solutions through a two-step equilibration process. The composition of each solution is described in Table 1. The first equilibration step used a total CPA concentration of 7.5% (v/v), while the second step used the same composition with the CPA concentration increased to 15% (v/v).

**Table 1:** Table 1. Composition and cooling characteristics of the cryopreservation protocols evaluated in this study. Concentrations of DMSO, ethylene glycol (EG), FBS, and basal medium (DMETC199) are expressed as % (v/v), whereas sucrose concentration is expressed in molarityComposition and steps of each protocol used in this study for 7.5% CPA concentration.

| Parameter | Slow freezing | Two-step freezing | Cold vitrification | Warm vitrification |
| --- | --- | --- | --- | --- |
| Initial temperature | 25 °C | 25 °C | 4 °C | 37 °C |
| Intermediate temperature | - | Approximately -10 °C | - | - |
| Final temperature | -80 °C | -196 °C | -196 °C | -196 °C |
| Approximate cooling time | >90 min | >20 min | <1 s for LN<sub>2</sub> immersion | <1 s for LN<sub>2</sub> immersion |
| Basal medium | - | DMETC199, 75% | DMETC199, 75% | DMETC199, 65% |
| FBS | 90% | 10% | 10% | 20% |
| DMSO | 10% | 7.5% | 7.5% | 7.5% |
| Ethylene glycol (EG) | - | 7.5% | 7.5% | 7.5% |
| Sucrose | - | - | - | 5 × 10<sup>-3</sup> M |

Accordingly, MDTs were first incubated for 10 min in an equilibration solution containing 7.5% DMSO and 7.5% ethylene glycol (v/v), followed by transfer for 3 min into a solution containing 15% DMSO and 15% ethylene glycol (v/v). Because the evaluated conditions were adapted from distinct preservation approaches, both the cooling profile and the complete cryopreservation formulation, including the FBS concentration and sucrose content, varied between conditions, as summarized in Table 1. Therefore, the experiments compared complete cryopreservation protocols rather than the effects of cooling profiles in isolation.

Three cooling strategies were evaluated:

1. **Warm vitrification:** Cryotubes were transferred directly from 37 °C into liquid nitrogen (−196 °C). Complete immersion of the cryotubes was achieved in <1 s.
2. **Cold vitrification:** Cryotubes containing approximately 1 mL of vitrification solution were placed horizontally in direct contact with dry ice in an insulated Styrofoam container for 1 min. Based on prior calibration, this exposure was expected to reduce the solution temperature to approximately 4 °C without inducing visible freezing, although the internal sample temperature was not directly measured during the present experiments. The cryotubes were then immediately immersed in liquid nitrogen (−196 °C), with complete immersion achieved in <1 s.
3. **Two-step freezing:** Following removal of most of the vitrification solution, several tens of microlitres were left surrounding the MDTs. Cryotubes were placed horizontally in direct contact with dry ice until the remaining solution visibly froze. Using long-handled forceps, the cryotubes were then fully immersed in liquid nitrogen for approximately 10 s, until vigorous boiling around the tubes had subsided.

The warm- and cold-vitrification conditions were designed to achieve rapid cooling without visible ice formation, whereas the two-step freezing condition intentionally permitted ice formation before liquid-nitrogen immersion and served as a comparative control. Samples from all three conditions were stored in the liquid phase of a liquid-nitrogen tank for one week.

Slow freezing was used as the comparative cryopreservation condition because it represents the conventional standard for preserving cell suspensions, allowing its suitability for three-dimensional MDTs to be evaluated against vitrification. Slow freezing was performed using a standard controlled-rate freezing protocol[28]. MDTs were suspended in freezing medium containing 10% DMSO and 90% FBS (v/v) and placed in a controlled-rate freezing container (5100-0001, Thermo Fisher Scientific), which provides a cooling rate of approximately −1 °C/min when stored at −80 °C. Samples were maintained at −80 °C for one week to match the storage period used for the other cryopreservation conditions.

For the warm-vitrification, cold-vitrification, and two-step-freezing conditions, cryoprotectant removal was performed using a stepwise dilution procedure immediately after warming. MDTs were transferred for 1 min into an unloading solution containing 15% DMSO and 15% ethylene glycol (v/v), followed by incubation for 5 min in a solution containing 7.5% DMSO and 7.5% ethylene glycol (v/v). The basal medium, FBS concentration, and presence of sucrose were matched to the formulation used for each cryopreservation condition, as detailed in Table 1. MDTs were then washed three times in pre-warmed culture medium for 5 min per wash at 37 °C to remove residual cryoprotective agents.

Each DMSO/ethylene glycol-based cryopreservation condition was evaluated without antioxidant supplementation and with the antioxidant concentrations described in Section 2.6. Following thawing and cryoprotectant removal, MDTs were reloaded into microfluidic chips and maintained under standard culture conditions for a four-day post-thaw recovery period. Fresh, non-cryopreserved MDTs were cultured in parallel for the same duration and served as time-matched controls. At the end of the recovery period, MDTs were fixed within the microfluidic devices and processed into FFPE blocks for subsequent histological and immunohistochemical analyses, as described in Section 2.7.

### 2.6 : Antioxidant supplementation

An antioxidant mixture (A3) composed of acetyl-L-carnitine (15 µM; A6706, Sigma-Aldrich, St. Louis, MO, USA), N-acetyl-L-cysteine (15 µM; A9165, Sigma-Aldrich), and α-lipoic acid (10 µM; T1395, Sigma-Aldrich) was prepared in PBS, aliquoted, and stored frozen until use. On the day of each experiment, aliquots were thawed and added to the cryoprotectant-containing equilibration and freezing solutions.

The antioxidants therefore remained present during cryoprotectant exposure, cooling, and cryogenic storage. However, they were not added to the cryoprotectant-unloading solutions or to the post-thaw culture medium, thereby limiting continued antioxidant exposure after thawing. In addition to the reference concentrations reported in the literature [25], 10-fold and 100-fold diluted formulations were evaluated to characterize the dose-dependent balance between cryoprotective activity and potential toxicity in three-dimensional MDTs.

The evaluated experimental arms were adapted from distinct cryopreservation approaches and therefore differed in both cooling profile and complete solution formulation. Consequently, they were compared as integrated cryopreservation protocols rather than as conditions designed to isolate the effect of a single parameter. Within each protocol, antioxidant supplementation was evaluated while the remaining cryopreservation conditions were kept unchanged.

### 2.7 : Tissue viability and immunohistochemistry

Tissue viability and structural integrity were assessed in cryopreserved MDTs and in fresh, non-cryopreserved MDTs cultured in parallel for matched durations. Sections from the original xenograft tumors and freshly microdissected MDTs fixed at day 0 were also included as baseline controls. At each experimental endpoint, MDTs cultured in the microfluidic devices were fixed and processed into FFPE blocks. Histological and immunostaining analyses were subsequently performed on tissue sections rather than directly within the microfluidic devices. Hematoxylin and eosin (H&E) staining was used to evaluate tissue morphology, including structural integrity, cellular organization, and the presence of necrotic regions. Proliferation, nuclear density, and apoptosis were assessed using Ki-67, DAPI, and cleaved caspase-3 (CC3), respectively.

Following fixation and paraffin embedding, FFPE blocks were sectioned at 4 µm thickness using a microtome and mounted on glass slides (TOM-1190, Matsunami). Slides were processed using the Discovery Ultra automated staining platform (Ventana Medical Systems, Roche, Switzerland).

After deparaffinization and rehydration, antigen retrieval was performed using Cell Conditioning #1 solution (Ventana, Tris-EDTA buffer, pH 7.8) for 60 min at 95 °C. Slides were incubated for 60 minutes at 37 °C with primary antibodies against Ki-67 (rabbit anti-Ki67, MA5-14520, Thermo Fisher Scientific) and cleaved caspase-3 (CC3, 9661S, New England Biolabs), both at a dilution of 1:600.

Detection was performed using the Discovery OmniMap HRP system (Ventana) with anti-rabbit secondary antibodies. Slides were counterstained with hematoxylin and bluing reagent (Ventana). Whole-slide images were acquired using an Aperio Verso 200 slide scanner (Leica Biosystems) equipped with a 20x objective (0.8 NA), corresponding to a resolution of 0.275 µm/pixel. Images were visualized using Aperio ImageScope software (Leica Biosystems).

Quantitative image analysis was performed using Visiomorph™ software (Visiopharm, Denmark). Marker levels were quantified using threshold-based segmentation, and results were expressed as the fraction of positively stained area relative to total tissue area.

### 2.8 : Statistical analysis

Statistical analyses were performed using GraphPad Prism (GraphPad Software, San Diego, CA, USA). Data are presented as mean ± SEM (standard error of the mean), with each point representing an individual MDT pooled across biological replicates. Group comparisons were performed using one-way ANOVA followed by Tukey’s multiple-comparisons test. All tests were two-sided, and p values < 0.05 were considered statistically significant.

## III. RESULTS

### 3.1 : Vitrification and slow freezing yield variable proliferation across cell lines

Warm vitrification was first compared to slow freezing using fresh MDTs to serve as comparative standard cryopreservation (Fig. 1). Following paraffin embedding and immunohistochemical analysis, proliferation (Ki-67) and apoptosis (CC3) marker levels were quantified for both cell lines. The two cell lines responded differently to cryopreservation. In the TOV112D model, vitrified MDTs showed slightly higher proliferation than slow-frozen samples, whereas in the 49F model, proliferation was slightly higher following slow freezing. In both cell lines, however, proliferation remained well below that of fresh controls (Fig. 1A). This effect was more pronounced in the 49F model, where overall proliferative activity was low across cryopreservation conditions. This is consistent with the already low baseline proliferation of fresh tissues in 49F. Apoptotic levels also varied depending on the condition and cell line (Fig. 1B). In the 49F model, vitrification resulted in a lower apoptotic area compared to slow freezing, whereas in the TOV112D model, apoptosis levels remained comparable between the two cryopreservation methods.

**Figure 1.**
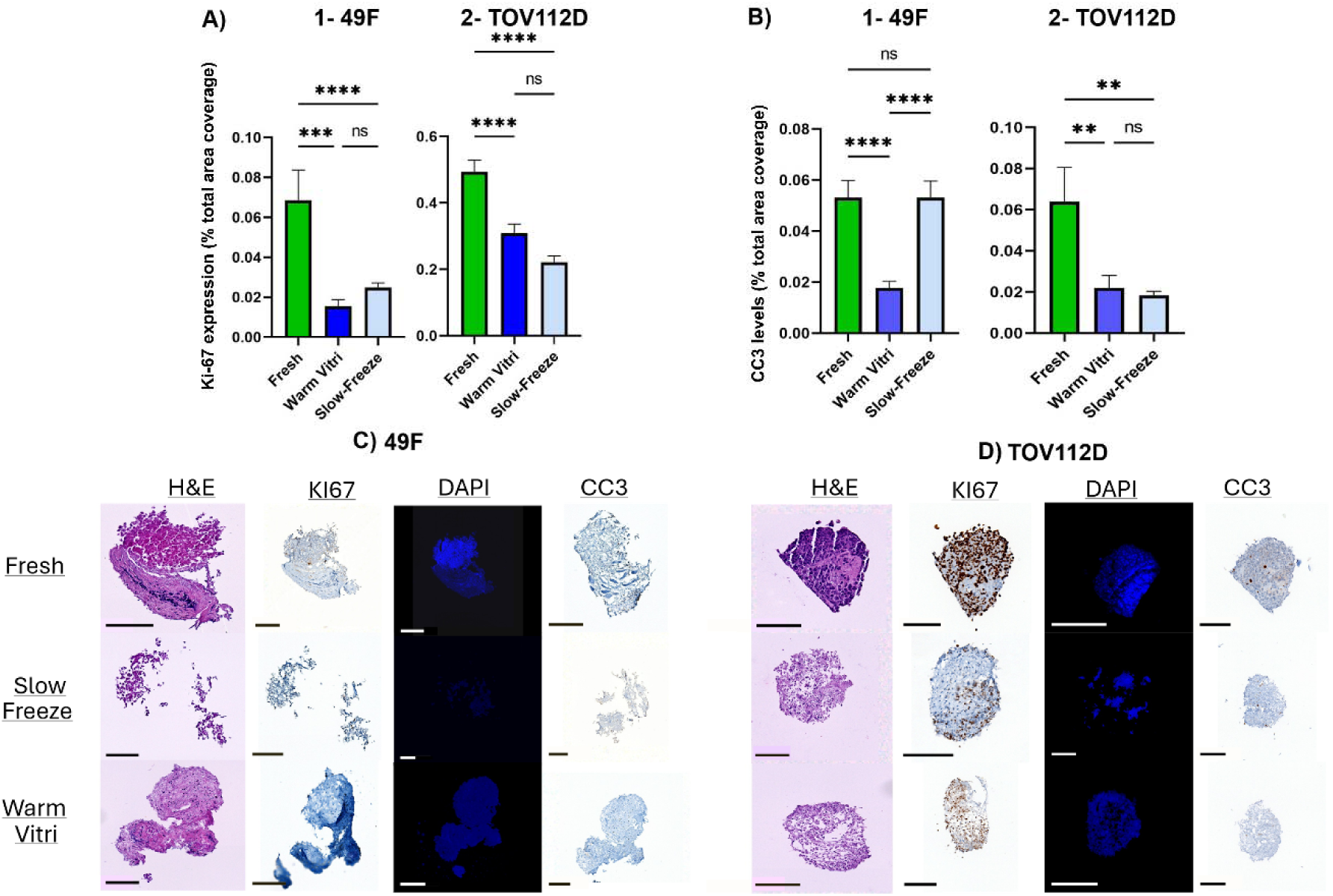
Comparison of slow freezing and warm vitrification for the cryopreservation of microdissected tumor explants (MDTs). (A) Ki-67-positive area and (B) CC3-positive area quantified in 49F (prostate) and TOV112D (ovarian) MDTs following fresh, slow-freezing, or warm vitrification conditions. (C–D) Representative H&E, Ki-67, DAPI, and CC3 staining of 49F and TOV112D MDTs under each cryopreservation condition. Data are presented as mean ± SEM (N = 3, n ≥ 31 for Ki-67; N = 3, n ≥ 40 for CC3). Statistical analysis was performed using one-way ANOVA followed by Tukey’s multiple-comparisons test. **P < 0.01, ***P < 0.001, ****P < 0.0001. Scale bars = 200 µm.

Histological analysis further illustrated differences in tissue integrity across conditions (Fig. 1C-D). H&E staining showed partial preservation of tissue architecture in both cryopreservation methods relative to fresh samples, while Ki-67, DAPI, and CC3 staining confirmed variability in proliferation, nuclear integrity, and apoptosis across conditions.

### 3.2 : Antioxidant supplementation increases proliferation, while temperature effects are cell line– dependent

Across all vitrification conditions, antioxidant supplementation with A3 increased proliferation compared to non-supplemented controls, as indicated by a higher proportion of Ki-67-positive areas (Fig. 2A). The highest proliferation levels were observed at the standard (1x) concentration, approaching those measured in fresh samples. Increasing A3 concentration beyond this level resulted in reduced proliferation, although values remained higher than without antioxidants. Morphological assessment and immunohistochemistry further showed reduced tissue integrity at higher A3 concentrations (Fig. 2C), consistent with decreased proliferation levels.

**Figure 2:**
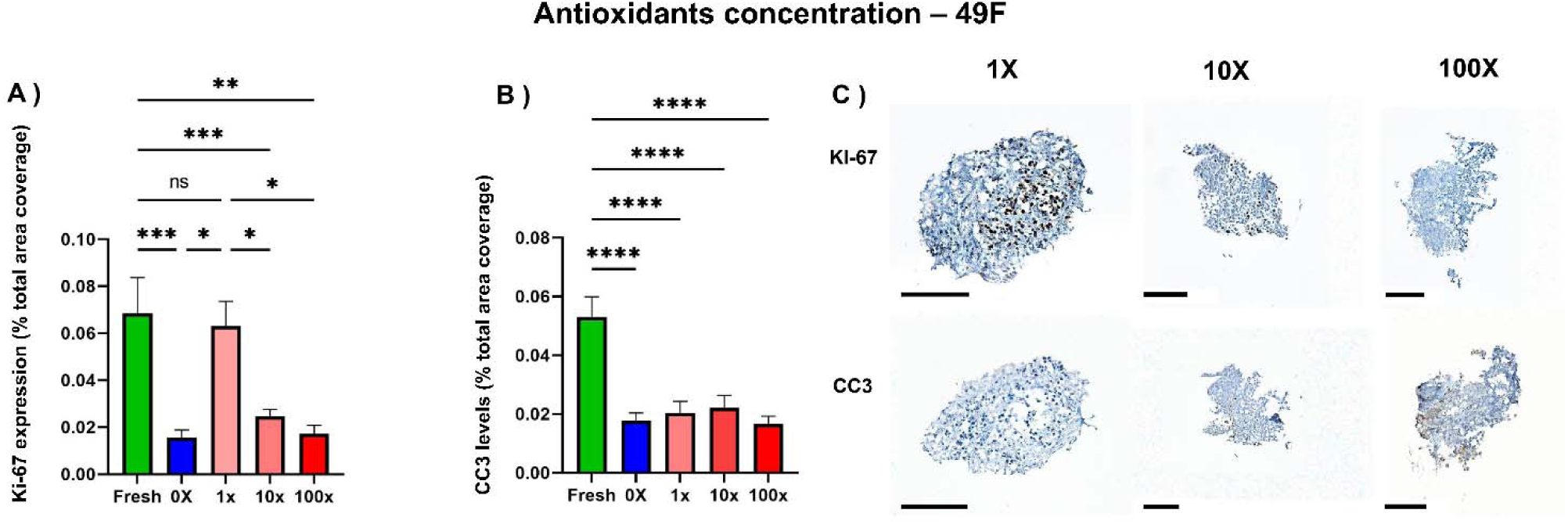
Impact of antioxidant concentration on the cryopreservation of microdissected tumor explants (MDTs). (A) and (B) show proliferative and apoptotic total areas, respectively, in the prostate cell line following cryopreservation with increasing concentrations of the antioxidant mix (A3). Proliferation levels comparable to fresh samples were observed at concentration 1X, while apoptotic levels remain. (C) shows proliferative staining (Ki-67) and apoptotic staining (CC3) for each of the increasing concentrations of A3). Data were analyzed using one-way ANOVA followed by Tukey’s multiple comparison test. *P < 0.05, **P < 0.01, ***P < 0.001, ****P < 0.0001. Scale bars represent 200 µm.

The effect of temperature prior to vitrification differed between cell lines. In the 49F model, pre-cooling to 4 °C (cold vitrification) resulted in higher proliferation compared to warm vitrification (37 °C) (Fig. 3A). In contrast, no significant difference between temperature conditions was observed in the TOV112D model (Fig. 3B). No MDTs could be recovered following vitrification initiated at −10 °C in the TOV112D model, and this condition was therefore excluded from analysis. Apoptotic levels remained comparable across antioxidant concentrations and temperature conditions (Fig. 2B, 3B).

**Figure 3.**
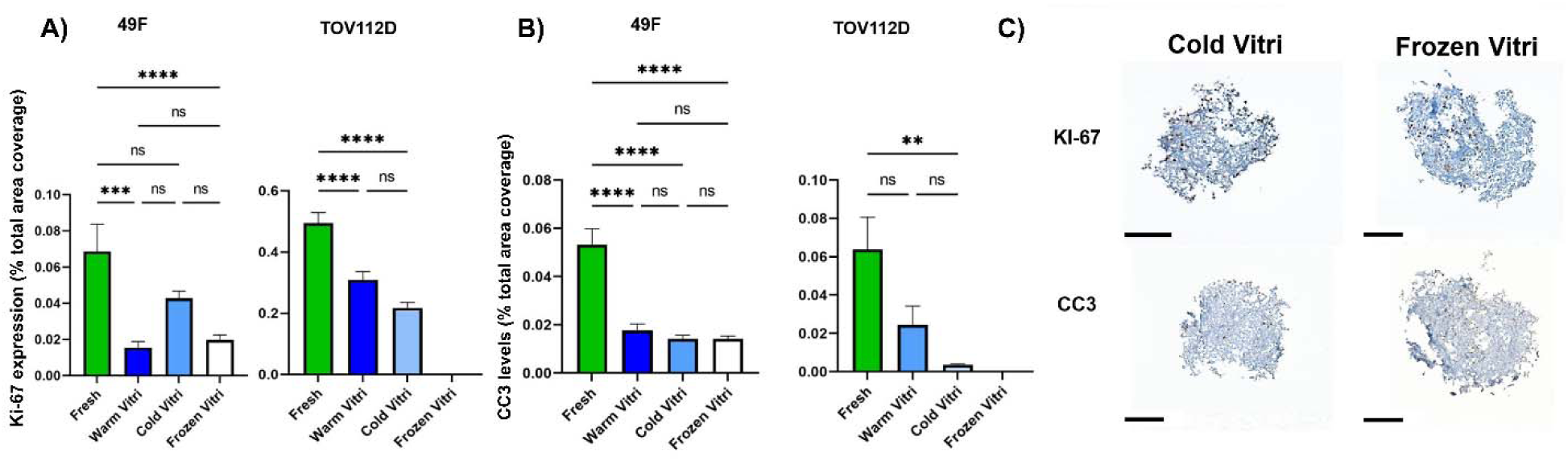
Effect of temperature gradient on the cryopreservation of microdissected tumor explants (MDTs). (A) Ki-67-positive area and (B) CC3-positive area quantified in 49F and TOV112D MDTs following vitrification under different starting temperature conditions (−10 °C, 4 °C, and 37 °C). No viable TOV112D MDTs were recovered following vitrification initiated at −10 °C. (C) Representative Ki-67 and CC3 staining of MDTs following vitrification under the different temperature conditions (representative images for warm vitrification are shown in *Fig. 1*). Data are presented as mean ± SEM. Statistical analysis was performed using one-way ANOVA followed by Tukey’s multiple-comparisons test. *P < 0.05, **P < 0.01, ***P < 0.001, ****P < 0.0001. Scale bars = 200 µm.

### 3.3 : Combined vitrification parameters restore proliferation close to fresh levels

Based on the results from Sections 3.1 and 3.2, a combined cryopreservation protocol was defined using the warm vitrification protocol (37 °C to -196 °C), A3 antioxidant supplementation (1x concentration), and standard CPA conditions. This protocol was selected for its consistently higher proliferation outcomes across experiments. Proliferation levels following this combined protocol approached those of fresh samples in both cell lines (Fig. 4A). In the 49F model, vitrified MDTs reached 98% of the proliferative area measured in fresh samples, while in the TOV112D model, proliferation reached 69% of fresh levels. In both cases, these values were higher than those observed in previous vitrification conditions.

**Figure 4:**
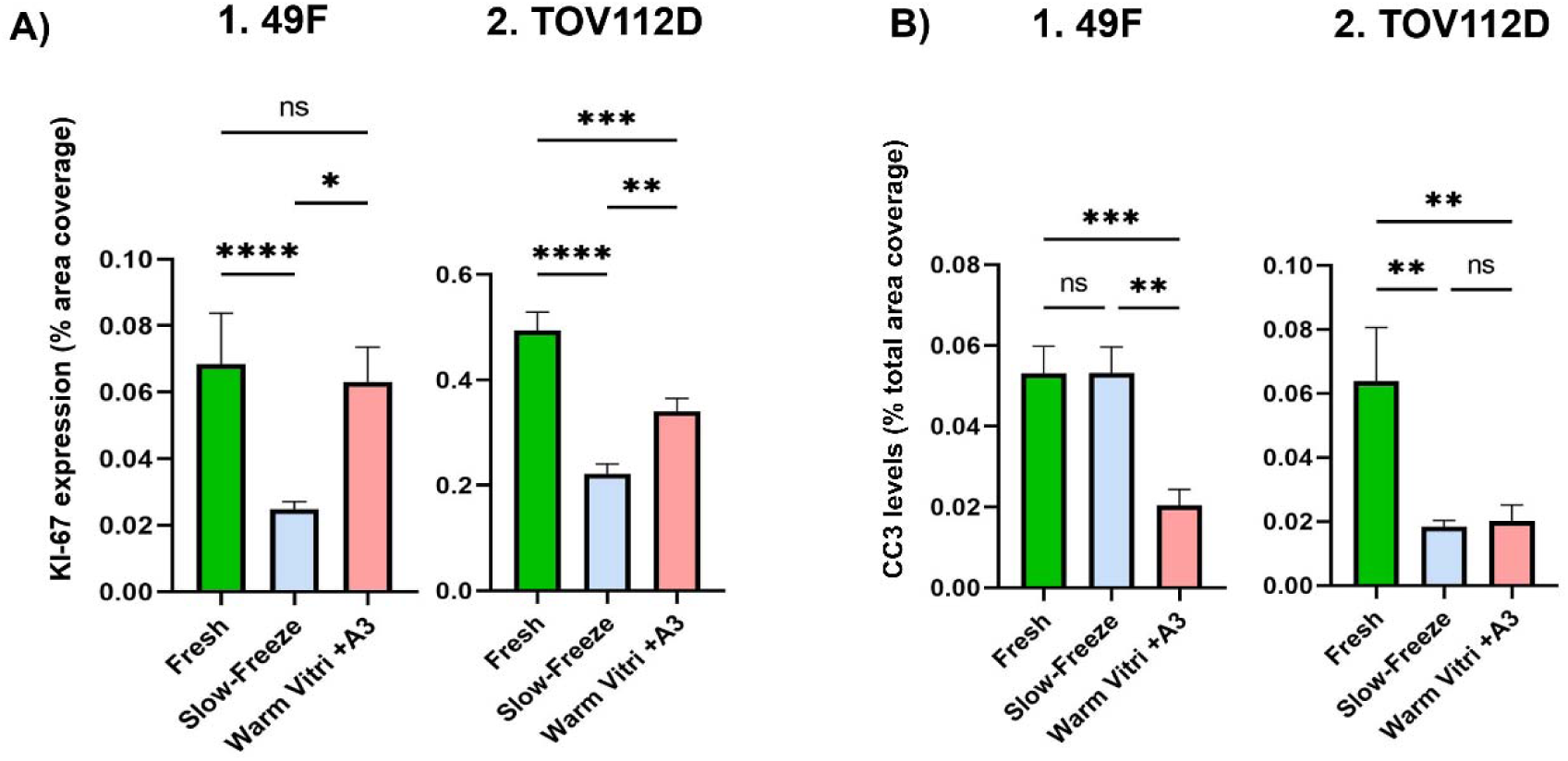
Optimized protocol with solution composition, temperature gradient and A3 concentration. A.1) Proliferative total area in the prostate cell line, showing no significant difference between fresh and vitrified samples.) A.2) Proliferative total area in the ovarian cell line, showing a significant increase in signal for vitrified samples compared to the slow-freezing technique. (N =3, n ≥24) B.1) Apoptotic total area in the prostate cell line, showing a significantly reduced area of dying cells in vitrified conditions. B.2) Apoptotic total area in the ovarian cell line, highlighting significantly reduced apoptosis levels for both cryopreservation methods compared to fresh. Representative stainings are presented in Fig. 1 and 2 (N =3, n ≥44). Data were subjected to a one-way ANOVA followed by Tukey’s multiple comparison test. *P<0.05, **P<0.01, ***P<0.001, ****P<0.0001.

Apoptotic levels remained low following vitrification in both cell lines (Fig. 4B). In the 49F model, apoptosis was significantly reduced compared to slow freezing, while in the TOV112D model, both cryopreservation methods resulted in lower apoptosis compared to fresh samples. In contrast, slow freezing combined with antioxidant supplementation did not result in increased proliferation compared to previous conditions, with proliferative levels remaining comparable to those observed without A3.

### 3.4 : PRE-CRYOPRESERVATION CULTURE IMPROVES POST-THAW PROLIFERATION IN 49F MDTs

To isolate the effect of pre-cryopreservation culture, this experiment was performed in 49F MDTs using vitrification without antioxidant supplementation. MDTs were maintained in culture for 24 or 48 h before vitrification using either microfluidic chips or standard Petri dishes (Fig. 5). Both recovery durations increased post-thaw proliferation compared with MDTs vitrified immediately after microdissection, with no additional benefit observed between 24 and 48 h of recovery (Fig. 5A).

**Figure 5.**
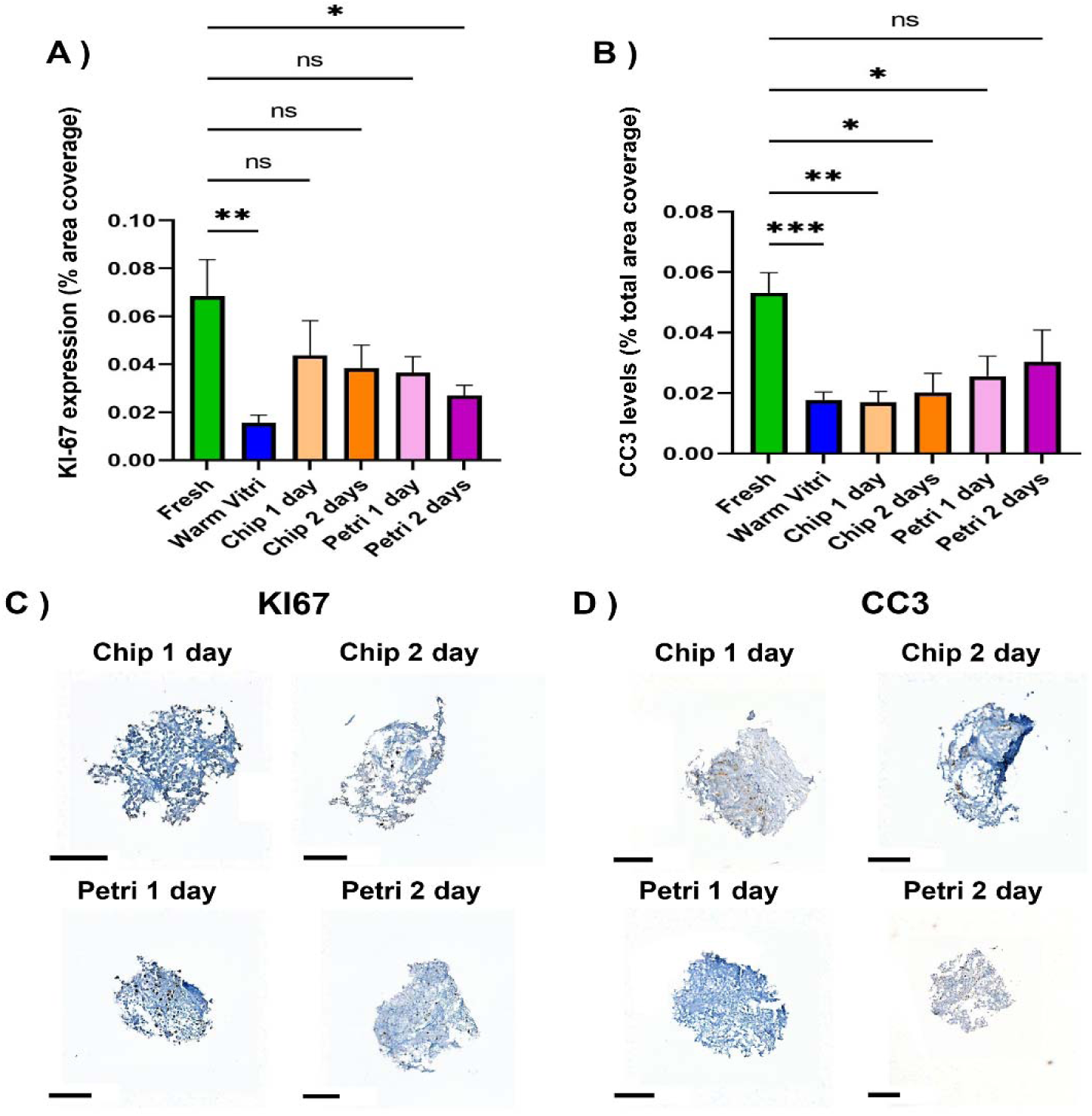
Effect of pre-cryopreservation recovery culture on the viability of microdissected tumor explants (MDTs). (A) Ki-67-positive area and (B) CC3-positive area quantified in 49F MDTs following vitrification with no recovery period or following 1 or 2 days of recovery culture in either standard Petri dishes or microfluidic chips prior to cryopreservation. (C) Representative Ki-67 staining and (D) representative CC3 staining of MDTs following recovery culture under each condition. Data are presented as mean ± SEM (N = 3, n ≥ 22 for Ki-67; N = 3, n ≥ 16 for CC3). Statistical analysis was performed using one-way ANOVA followed by Tukey’s multiple-comparisons test. *P < 0.05, **P < 0.01, ***P < 0.001. Scale bars = 200 µm.

No significant differences in proliferative area were observed between MDTs recovered in microfluidic chips and those maintained in Petri dishes. Apoptotic levels remained low and were not significantly affected by the presence, duration, or culture platform of the recovery period (Fig. 5B). Representative Ki-67 and CC3 staining are shown in Fig. 5C-D. Because this experiment was limited to the 49F model and was conducted without antioxidant supplementation, the interaction between recovery culture, A3 supplementation, and tumor model was not assessed.

## IV. Discussion

This work systematically explored cryopreservation strategies for microdissected tumor tissues (MDTs), addressing a major limitation in the use of complex *ex vivo* tumor models: the lack of reliable and reproducible preservation methods. While slow freezing remains the most commonly used approach for cells, this method remains ill-adapted to complex 3D models. Our results demonstrate that vitrification yields superior preservation of tissue structure and viability in MDTs, provided that key parameters are carefully optimized.

Previous studies on tumor cryopreservation have typically focused on isolated parameters (such as cryoprotectant composition) or on a single cancer model and often rely on technically complex approaches such as nanoparticle-assisted vitrification [29], [30], [31], [32]. Many studies assess tissue viability immediately after thawing, without allowing for post-thaw recovery, which limits their relevance for functional assays. In contrast, this study systematically evaluates multiple parameters, including temperature, antioxidant supplementation, and recovery conditions, across two distinct cancer models, using a microfluidic chip that enables controlled culture and longitudinal assessment. This approach highlights not only the potential of vitrification, but also the extent to which the outcome depends on protocol design and biological context.

Across the evaluated staining modalities, vitrification generally resulted in better preservation of tissue morphology than slow freezing, as evidenced by H&E and DAPI staining. However, the apoptotic response to cryopreservation differed substantially between the two tumor models. In 49F MDTs, slow freezing produced greater CC3-positive areas than vitrification, whereas a comparable increase was not observed in TOV112D MDTs. Moreover, the relatively low apoptotic signal measured in some cryopreserved TOV112D conditions did not correspond to complete preservation of proliferative capacity. These findings indicate that cryopreservation outcomes cannot be inferred from apoptosis or structural integrity alone and that Ki-67, CC3, and tissue morphology must be interpreted together. They also suggest that susceptibility to cryopreservation-induced injury depends on intrinsic characteristics of the tumor model, including baseline proliferative activity, tissue organization, and stress sensitivity.

Antioxidant supplementation with A3 improved post-thaw proliferation under vitrification conditions, particularly at concentrations consistent with prior literature [25]. However, higher concentrations led to diminished benefits and signs of toxicity, highlighting a narrow effective concentration range. In contrast, A3 supplementation did not produce a comparable improvement following slow freezing. This suggests that oxidative stress contributes to vitrification-associated injury but may not be the principal limitation of conventional slow freezing in three-dimensional MDTs. Damage arising from ice formation, prolonged cooling, or nonuniform cryoprotectant distribution within the tissue may remain dominant under slow-freezing conditions and may not be corrected by antioxidant supplementation alone. Antioxidant-based protection must therefore be optimized in the context of the complete cryopreservation protocol rather than considered independently.

The cooling protocol preceding liquid-nitrogen immersion also influenced post-thaw outcomes in a cell line-dependent manner. In the 49F model, the cold-vitrification protocol produced higher proliferation than warm vitrification, whereas no significant difference was observed in the TOV112D model. Because these conditions differed not only in their initial cooling procedure but also in aspects of solution composition and retained volume, the observed effects cannot be attributed to temperature alone. The results nevertheless indicate that the conditions immediately preceding vitrification can influence tissue recovery and may interact with model-specific biological characteristics, such as cold-stress responses, membrane behavior, and intrinsic sensitivity to cryopreservation.

By integrating the evaluated parameters, we identified the best-performing vitrification protocol under the conditions tested, combining warm vitrification (37 °C to −196 °C), controlled cryoprotectant exposure, and A3 supplementation at the reference concentration. Under these conditions, post-thaw proliferation reached 98% and 69% of fresh-control levels in the 49F and TOV112D models, respectively. Although these findings demonstrate that high levels of functional preservation are achievable, the difference between the two models highlights an important challenge for application to freshly collected tumor specimens, for which extensive sample-specific optimization would not be practical. Further experiments across a broader range of tumor models and patient-derived tissues will therefore be required to identify parameters that produce more consistent outcomes despite variations in tissue composition, metabolic state, and intrinsic stress sensitivity. Such studies may support the development of either a broadly applicable protocol or a limited set of preservation conditions selected according to measurable tissue characteristics.

The present findings also place tumor cryopreservation in the broader context of vitrification strategies successfully implemented in assisted reproductive technologies. Vitrification has become the preferred approach for the preservation of oocytes and embryos because it minimizes ice crystal formation and enables high post-thaw survival and developmental competence. While these successes demonstrate the ability of vitrification to preserve complex multicellular systems, our results indicate that direct translation to tumor explants is not straightforward. Unlike embryos, which are highly organized and relatively homogeneous structures, tumor explants comprise heterogeneous populations of malignant, stromal, immune, and vascular cells embedded within a complex extracellular matrix. These features are likely to influence cryoprotectant diffusion, osmotic responses, and susceptibility to cryoinjury, emphasizing the need for tissue-specific optimization of vitrification protocols.

Importantly, a short recovery period following microdissection improved post-thaw proliferation in 49F MDTs vitrified without antioxidant supplementation. A 24-h recovery period was sufficient, with no additional benefit observed after 48 h. This finding suggests that mechanical stress induced during tissue preparation contributes to cryopreservation sensitivity and that allowing the tissue to stabilize before freezing may improve its subsequent recovery. The effect was comparable in microfluidic chips and conventional Petri dishes, indicating that this step can be implemented without specialized culture equipment. However, because the experiment was performed in only one tumor model and without A3 supplementation, it remains unclear whether pre-cryopreservation recovery provides an additive benefit when combined with the optimized antioxidant-containing protocol or whether the effect generalizes to other tumor types.

Taken together, these results position vitrification as a promising strategy for preserving complex tumor explants, provided that both physical and biological parameters are considered. Compared to simpler in vitro models such as cell suspensions or organoids, MDTs retain key features of tumor architecture and heterogeneity, making their preservation particularly valuable for translational applications. While current cryopreservation approaches for such models remain inconsistent and often poorly standardized, our work provides a framework for rational protocol optimization, balancing reproducibility with adaptability.

This study was limited to murine tumor xenografts from two cancer models, and validation in additional tumor types, particularly patient-derived tissues, will be required to establish broader applicability. Long-term storage beyond one week and extended post-thaw culture were also not investigated, leaving open questions regarding preservation durability and long-term tissue function. Moreover, the evaluated conditions represent only a subset of the cryopreservation strategies described in the literature. Additional parameters, including alternative cryoprotectant formulations, loading and unloading procedures, cooling and warming profiles, and preservation solutions adapted from organ transplantation, could further improve tissue recovery. Because systematically testing every possible combination would be impractical, future studies should use staged experimental designs to identify the parameters with the greatest influence on preservation outcomes. The central challenge will be to develop the simplest and most robust protocol capable of producing consistent results across heterogeneous tumor tissues. Investigating the mechanisms underlying differential responses to temperature, antioxidants, and solution composition may also support more rational protocol selection and reduce the need for extensive sample-specific optimization.

Although only xenograft-derived MDTs were evaluated in this study, the proposed workflow is directly compatible with patient-derived explants generated using existing microdissection protocols. Patient samples are expected to exhibit greater inter-patient variability in cellular composition, stromal content, and treatment history, which may influence cryopreservation outcomes. However, the systematic optimization strategy presented here provides a framework that can readily be adapted to these more heterogeneous clinical specimens. Successful translation to patient-derived tissue could facilitate centralized biobanking, improve logistical flexibility around surgical specimen collection, and enable broader implementation of explant-based functional assays for precision oncology.

## V. Conclusion

While vitrification offers clear advantages over conventional slow-freezing methods for tissue cryopreservation, our results demonstrate that its performance is highly dependent on protocol design and biological context. Rather than being universally superior, vitrification requires careful optimization of key parameters to achieve reliable outcomes in complex tissue models.

By systematically evaluating the effects of antioxidant supplementation, initial temperature, and post-microdissection recovery, we identified a combination of conditions that consistently improves the preservation of viability, proliferation, and morphology in microdissected tumor explants. These findings highlight that successful cryopreservation of MDTs depends not only on physical parameters such as cooling rate, but also on biological factors including tissue stress and recovery capacity.

Although further validation is required across additional tumor types, patient-derived samples, and longer storage durations, this work establishes a rational framework for the optimization of vitrification protocols in complex tissue systems. Ultimately, enabling reliable cryopreservation of MDTs could facilitate centralized biobanking, improve experimental reproducibility, and accelerate the integration of explant-based models into precision oncology workflows.

## VI. CRediT authorship contribution statement

**Tommy Brasseur:** Conceptualization, Data curation, Formal analysis, Investigation, Methodology, Software, Validation, Visualization, Writing – original draft, Writing – review & editing.

**Gabriel Pagé:** Conceptualization, Data curation, Formal analysis, Investigation, Methodology, Software, Validation, Visualization, Writing – review & editing.

**Amélie St-Georges-Robillard:** Investigation, Methodology, Project administration, Visualization, Writing – review & editing.

**Anne-Marie Mes-Masson:** Funding acquisition, Investigation, Project administration, Resources, Supervision, Writing – review & editing.

**Thomas Gervais:** Conceptualization, Funding acquisition, Investigation, Methodology, Project administration, Resources, Supervision, Writing – review & editing.

## VII Acknowledgements

We would like to thank the Microfluidics Core Facility team, Jennifer Kendall-Dupont, and Benjamin Péant, for their valuable support with MDT generation and culture, as well as Kim Leclerc-Desaulniers for assistance with animal handling. We are also grateful to the Molecular Pathology Core Facility, particularly Véronique Barrès, for IHC and IF support. We want to thank Fayrouz Annab and Dr. Fred Saad for providing the prostatic cell line. This work was funded by the Natural Sciences and Engineering Research Council of Canada (NSERC) and the Programme de Soutien aux Organismes (PSO) [VAL1687-#53688] from the Ministère de l’Économie et de l’Innovation et de l’Énergie du Québec. This research was carried out in part using the Microfluidics Core Facility, which is supported by a Terry Fox Research Institute New Frontiers Program Project Grant #1123, and the TransMedTech Institute and its main financial partner, the Canada First Research Excellence Fund. We also thank Fertilys Inc. for fruitful discussions and for sharing their cryopreservation protocol.

## IX. Declaration of generative AI and AI-assisted technologies in the manuscript preparation process

During the preparation of this work, OpenAI ChatGPT was used during late stages of manuscript preparation to assist with language refinement, formatting consistency, and improvement of grammar and readability. This AI technology was not used to generate new scientific content, perform data analysis, nor propose original ideas. All scientific content, interpretations, and conclusions were conceived and written by the authors. After using this tool, the authors reviewed and edited the content as needed and took full responsibility for the content of the published article.

