## Supplementary figures and images for "Enhancing the cryopreservation of *ex vivo* 3D tumor models using vitrification strategies"

### Supplemental Figure 1

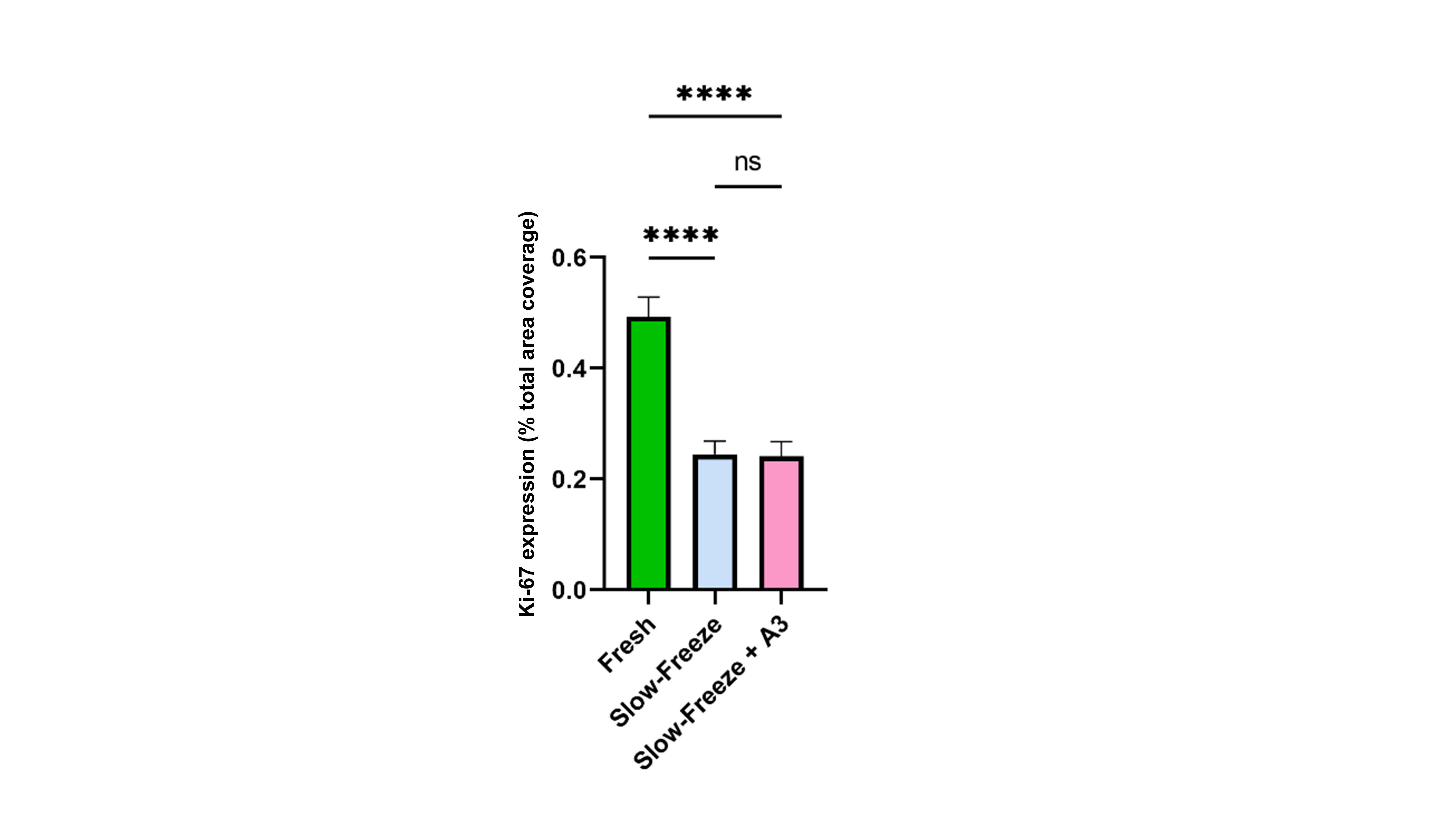
